# Glu-370 in the Large Subunit Influences the Substrate Binding, Allosteric, and Heat Stability Properties of Potato ADP-glucose Pyrophosphorylase

**DOI:** 10.1101/054106

**Authors:** Ayse Bengisu Seferoglu, Seref Gul, Ugur Meric Dikbas, Ibrahim Baris, Kaan Koper, Mahmut Caliskan, Gul Cevahir, Ibrahim Kavakli

## Abstract

ADP-glucose pyrophosphorylase (AGPase) is a key allosteric enzyme in plant starch biosynthesis. Plant AGPase is a heterotetrameric enzyme that consists of large (LS) and small subunits (SS), which are encoded by two different genes. In this study, we showed that the conversion of Glu to Gly at position 370 in the LS of AGPase alters the heterotetrameric stability along with the binding properties of substrate and effectors of the enzyme. Kinetic analyses revealed that the affinity of the LS^E370G^SS^WT^ AGPase for glucose-1-phosphate is 3-fold less than for wild type (WT) AGPase. Additionally, the LS^E370G^SS^WT^ AGPase requires 3-fold more 3-phosphogyceric acid to be activated. Finally, the LS^E370G^SS^WT^ AGPase is less heat stable compared with the WT AGPase. Computational analysis of the mutant Gly-370 in the 3D modeled LS AGPase showed that this residue changes charge distribution of the surface and thus affect stability of the LS AGPase and overall heat stability of the heterotetrameric AGPase. In summary, our results show that LS^E370^ intricately modulate the heat stability and enzymatic activity of the AGPase.

**Abbreviations**
AGPase: ADP-glucose pyrophosphorylase
BSA: bovine serum albumin
DTT: dithiothreitol
G1P: glucose-1-phosphate
IPTG: isopropyl-P-D-thiogalactopyranoside
LS: large subunit
3PGA: 3-phosphoglyceric acid
Pi: inorganic phosphate
SS: small subunit
TBS: Tris-buffered saline
WT: wild type

## 1. Introduction

Starch is a staple in the diet of much of the world’s population and is also widely used in different industries as a raw material [1]. The first committed step of starch biosynthesis in plants is catalyzed by ADP-glucose pyrophosphorylase (AGPase, EC 2.7.7.27). AGPase, an allosteric enzyme, catalyzes the rate limiting reversible reaction and controls the carbon-flux in the ɑ-glucan pathway by converting Glucose-1-phosphate (G1P) and ATP to ADP-glucose and pyrophosphate using Mg^2+^ as the cofactor [1–3]. Regulation of almost all AGPases depends on the 3-phosphoglyceric acid to inorganic phosphate ratio (3PGA/Pi). While 3PGA functions as the main stimulator, P_i_ inhibits the activity of enzyme [4–6]. Plant AGPases consist of pairs of small (SS, or α) and large (LS, or β) subunits, encoded by two distinct genes, thereby constituting a heterotetrameric structure (α_2_β_2_) [7]. In potato tuber AGPase, the sequence identity between the LS and the SS is 53% suggesting a common ancestral gene [8, 9]. The molecular mass of tetrameric AGPases ranges from 200 to 240 kDa depending on the tissue and plant species. Specifically, molecular masses of LS and SS in potato tuber AGPase are 51 kDa and 50 kDa, respectively [10].

When expressed in bacteria, the SS of potato tuber AGPase can form a catalytically active homotetrameric enzyme with defective allosteric properties [11, 12]. The LS was incapable of self-assembly and therefore was assumed to be essential for modulating the allosteric properties of the active heterotetrameric enzyme [13]. Evidence indicates that the LS may bind to the G1P and ATP substrates [14–16]. The binding of the LS to its substrates may allow the LS to interact cooperatively with the SS for binding substrates and effectors, which, in turn, influences net catalysis [14, 15,17,18].

Manipulation of the AGPase activity and its heat stability gets much attention in increasing the starch yield of plants because it catalyzes the first committed step as well as the rate limiting step of starch biosynthesis [19–22]. Modulating the heat stability is especially important for cereal AGPases because elevated temperature is a major environmental factor that greatly reduces grain yield and seed weight [23, 24]. Among AGPases, the potato tuber AGPase is much more heat stable than many others [10, 23]. It has been shown that the lack of a QTCL motif, which is found on the SS of potato tuber AGPase, is one of the reasons for the heat labile profile of the maize AGPase [25]. Thermal stability of the enzyme has been shown to not only depend on the robustness of the LS-SS interactions but also on the binding of the effector molecules to the allosteric sites [23, 26, 27].

The structure of homotetrameric potato SS [28] and *A.tumefaciens* [29] AGPases were solved by X-ray crystallography. Neither the LS nor the heterotetrameric AGPase (α_2_β_2_) structure have been solved yet. It is, therefore, homology-modeling approach was taken to understand structure-functions of heterotetrameric plant AGPases [30–34]. The modeling studies enable us to identify critical amino acid residues at the interface of the AGPase that are important for the LS-SS interaction and allosteric regulations [30–34]. Furthermore, based on the described heterotetrameric AGPase assembly model, LS-SS dimers are formed by side-by-side interactions and then LS-SS dimers interact in a head to tail orientation to form heterotetrameric AGPases [34]. Although there are several studies showed the importance of heterotetrameric assemblies in enzyme functions [13, 35–37], amino acids that contribute structural stability of heterotetrameric AGPase is not well defined. In this study, we demonstrated that an amino acid (LS^E370^), close to the LS interface, is important for maintaining the heat stability of the heterotetrameric structure of the AGPase. Computational analysis on the 3D model of LS AGPase revealed that LS^E370G^ mutation disrupts hydrogen bond interaction network and electrostatic interactions in that region. In addition, biochemical characterization showed that LS^E370G^ SS^WT^ AGPase was less heat stable and showed altered affinities towards G1P and 3PGA. Our data demonstrate Glu-370 is critical for the structural stability of the LS subunit itself and the overall heterotetrameric AGPase, and influences the allosteric, substrate binding and heat stability properties of the AGPase.

## 2. Materials and Method

### 2.1 Site-directed mutagenesis

Glutamate was replaced into glycine at the position of 370 in the potato LS AGPase cDNA using PCR with appropriate primers (Table S1) as previously described [38]. The PCR products were digested with DpnI to remove template plasmid DNA and transformed into *E. coli* DH5α. The presences of the mutations were confirmed by DNA sequencing (Macrogen Inc, Netherlands).

### 2.2 Iodine staining

Iodine staining was performed by exposing *E.coli* AC70R1-504 (glgC^−^) cells containing wild type and mutant AGPase to iodine vapor after cells were grown overnight on Kornberg’s medium (1.1% K_2_HPO_4_, 0.85% KH_2_PO_4_, 0.6% yeast extract, 1.5% agar, pH7.0) supplemented with 1% glucose, 50 μg/ml kanamycin, and 50 μg/ml spectinomycin.

### 2.3 Protein expression

The wild type and LS^E370G^ AGPases were expressed in *E.coli* AC70R1-504 (glgC^−^) cells as previously described [38]. *E.coli* AC70R1-504 cells were transformed with the plasmids pML7 and pML10 containing potato LS and SS cDNAs, respectively. Cells were grown in LB medium containing 50 μg/ml spectinomycin and 50 μg/ml kanamycin. Once absorbance reached 1.0-1.2 at OD_600_, the cells were induced with 0.2 mM IPTG and 10 mg/L nalidixic acid at room temperature for 20 h.

### 2.4 Native PAGE analysis

The heterotetrameric assembly properties of the LS^E370G^ mutant AGPase with wild type SS were investigated using 3-13% gradient native polyacrylamide gels followed by Western blot using potato anti-LS and anti-SS antibodies as previously described [27].

### 2.5 Heat stability

Cells, expressing mutant and wild type LSs in pML7 plasmids and wild type SS in pML10 plasmid, were harvested in buffer containing 50 mM HEPES at a pH of 7.5, 5 mM phosphate buffer (K_2_HPO_4_/KH_2_PO_4_) 5 mM MgCl_2_ and 10% glycerol. Aliquots of cell-free extracts of the wild type and mutant AGPases were placed in a water bath at 65°C for 5 and 10 min and then cooled on ice. All preparations were subsequently analyzed with native page followed by Western blot as described above using 0.5 μg, 1 μg and 2 μg of total protein.

AGPase activities in the crude cell extracts were also measured using forward direction assay, as described in the kinetic characterization part, in the presence of saturated substrates (5 mM) and the 3PGA concentration (10mM). Blank samples were complete reaction mixtures without enzyme. To get measurable activity a total of 10 μg of protein were used in the assay. Duplicate samples were left on ice and their activity was taken to be 100%.

### 2.6 His-tagged protein expression and purification

The recombinant polyhistidine-tagged wild type and mutant AGPases were expressed and purified as described herein. Plasmids pSH228 and pSH275 containing the AGPase SS and LS coding sequences [39], respectively, were sequentially transformed into *E.coli* EA345 cells lacking endogenous AGPase activity due to the null mutation in its structural gene *glgC* [18]. Three colonies were inoculated into 25 ml of Luria Broth (LB) medium containing 0.4% glucose, 100 μg/ml ampicillin, and 50 μg/ml kanamycin. The starter culture was grown overnight at 37°C and transferred to 1 L of LB medium. When the OD_600_ of the culture was reached 1.0, the cells were induced with 0.1 mM IPTG, and the protein expression was induced for 18 h at room temperature. The cells were harvested and disrupted by sonication in 25 ml of binding buffer (25 mM HEPES-NaOH, pH8.0, 5% glycerol) supplemented with 0.5 mg/L lysozyme, EDTA-free protease inhibitor cocktail (Sigma), and 1 mM PMSF [40]. The soluble fraction was separated by centrifugation and passed through Macroprep DEAE resin (BIORAD). After extensive washing in binding buffer, AGPase was eluted with binding buffer containing 0.3 M NaCl and then loaded onto an immobilized metal affinity column (TALON metal affinity resin, Clontech). The column was washed with binding buffer containing 0.3 M NaCl and 5 mM imidazole. The protein was eluted with a 5-100 mM imidazole gradient in binding buffer containing 0.3 M NaCl. Fractions containing AGPase activity were combined and concentrated using 30 kDa cut-off Amicon ultra centrifugal filters (Millipore). The concentrated protein was divided into small aliquots and stored at −80°C until use.

### 2.7 Kinetic characterization

A non-radioactive endpoint assay was used to determine the amount of PPi produced by monitoring the decrease in the NADH concentration. The standard reaction mixture contained 50 mM HEPES-NaOH at a pH7.4, 15 mM MgCl_2_, 5.0 mM ATP, 5.0 mM G1P, and 3 mM DTT in a total volume of 100 μl. The assays were initiated by the addition of the enzyme, and the reactions were performed in a water bath at 37°C followed by termination by boiling for 2 min. AGPase activities were linear with respect to time and the amount of enzyme. The reactions were developed by adding 150 μl of coupling reagent to each tube, and the A_340_ was determined using a 96-well microplate reader. The coupling reagent was prepared as described in [41]. with minor modifications. All enzymes used in the coupling reagent were purchased from Sigma (Dorset USA). The coupling reagent contained final concentrations of 25 mM imidazole, 4.0 mM MgCl_2_, 1.0 mM EDTA, 0.2 mM NADH, 1.0 mM F6P, 0.005 units of PP_i_-dependent phosphofructokinase, 0.1 units of aldolase, 0.2 units of triose phosphate isomerase, and 0.3 units of glycerophosphate dehydrogenase per reaction at a pH7.4. PPi production was determined from a standard curve using increasing amounts of PP_i_ in complete reaction mixtures without enzyme.

The K_m_ and A_0.5_ values were determined in reaction mixtures in which one substrate or effector was added in varying amounts, and the other reaction components were saturated (5 mM). Kinetic constants were calculated by non-linear regression analysis using Prism software (Graph Pad). I_0.5_ values for P_i_ were determined in the presence of 1.0 and 2.5 mM 3PGA by adding increasing amounts of P_i_. Students’ t-test was used.

### 2.8 Reducing and non-reducing conditions for AGPase

Cell-free extracts were prepared as described previously [26]. For reducing conditions, samples were heated at 95 °C for 5 min with Laemmli sample buffer containing 4 mM DTT. For non-reducing conditions, Laemmli sample buffer without any reducing agent was used. The samples were exposed to 10% SDS–polyacrylamide gels followed by Western blotting.

### 2.9 Minimizing the Energy of AGPase structure

3D structures of large subunit and heterotetrameric assembly of AGPase modeled by our group previously [34] were used for computational analysis. Using the NAMD (v. 2.6) [42] and VMD (v. 1.9.1) program packages structure of hetero tetrameric AGPase was solvated in a rectangular box with TIP3P water molecules having a minimum of 10Å distance from the closest atom of the protein to the boundary, and then counter ions were added to neutralize the system. Initially only the side chains were minimized for 10000 steps. Subsequently all atoms were minimized for 10000 steps without pressure control. Then the system was heated up to 310K by increasing the temperature 10K and 10ps simulation was performed at each step.

### 2.10 Protein Stability Calculations

FoldX uses statistical energy terms, structural descriptors and an empirical potential obtained from addition of weighted physical energy terms (e.g. van der Waals interactions, solvation, hydrogen bonding, and electrostatics) to predict protein energetics and to provide quantitative estimation of intermolecular interactions promoting the protein stability depending on the availability of 3D structures [43]. Furthermore FoldX can calculate the interaction energy between subunits of a protein complex via the structure based energy function. Previously it has been shown that FoldX is able to produce accurate predictions about the stability change upon point mutations [43] and interaction of globular proteins [44–46]. After minimizing the structures by following the procedure mentioned above, we used ‘Repair Object’ module of FoldX which rearranges the side chain positions but not backbone of the structure, to obtain better optimized and minimized structure. In all of the calculations structures minimized by NAMD and then ‘repaired’ by FoldX were used. To validate the stability data obtained from FoldX, we used I-Mutant 3.0 (structure mode) (http://gpcr2.biocomp.unibo.it/cgi/predictors/I-Mutant3.0/I-Mutant3.0.cgi) [47] and POPMUSIC server (https://soft.dezyme.com/home). To predict the effect of mutations on the Tm value of AGPase tetramer we used HOTMUSIC server. HOTMUSIC is very recently developed novel tool to predict thermal stability of proteins by predicting the change in Tm [48]. This tool uses standard and temperature dependent statistical potentials in combination with the neural network. Totally 1600 mutations with experimentally measured ΔTm data were used to obtain the parameters of the model [48].

## 3. Results and Discussion

### 3.1 The large subunit E370G mutation impairs glycogen production in a bacterial complementation assay

Modeling studies followed by mutagenesis of the potato AGPase identified a hotspot mutant AGPase for the interaction of head to tail LS-SS dimers, where an Arg residue at position 88 was replaced by Ala in the LS (LS^R88A^) [30,34]. In our previous study we employed an error prone-PCR random mutagenesis approach using the LS^A88^ AGPase cDNA as template to select second-site LS suppressor that could reverse the iodine staining deficiency of the *E.coli* glgC^−^ containing wild type SS AGPase cDNA [27]. Stained colonies were picked up and plasmids were isolated. To identify mutations each mutant were subjected to the sequencing as described in [27]. To see the effect of the second site revertants independently in AGPase function, plasmids containing mutant LS cDNAs were then purified and subjected to site-directed mutagenesis by PCR to convert the primary mutation Ala88 back to the corresponding wild type arginine residue.

The LS cDNAs containing only the secondary mutations were then co-expressed with wild type SS in *E.coli* glgC^−^ and their activity were assessed by iodine staining. This study yielded the following two group of mutants: one group of the LS mutants was able complement bacterial AGPase expressed with wild type SS cDNA in *E.coli* glgC^−^ and the other group of the LS mutants was not able to complement bacterial AGPase when the cells were exposed to the iodine vapor in *E.coli* glgC^−^. The first group of the LS AGPase mutants was characterized by biochemical methods and found to have enhanced heterotetrameric assemblies and comparable kinetic properties to wild type AGPase [27].

In this study, five revertants from the second group (RM4, RM5, RM25, RM27 and RM29) of the LSs were studied. We wished to identify the role of the interface amino acids of the LS in AGPase function we therefore mapped the position of each mutation on the modeled heterotetrameric AGPase. The results indicated that there are five mutations (LS^A113T^, LS^F123L^, LS^F324L^, LS^A113T^, and LS^E370E^) on the interface or close the interface of the LS AGPase. We then introduced these mutations on wild type LS cDNA by site directed mutagenesis. The LS cDNAs containing only these mutations were then co-expressed with the WT SS in *E. coli* glgC^−^, and their activity was assessed by iodine staining. The results indicated that the cells containing LS^A113T^, LSF^123L^, LS^F324L^, LS^A113T^, and WT SS stained as dark as cells that contained WT cDNAs of the LS and SS (data not shown) whereas the cells with LS^E370G^ displayed no iodine staining phenotype compared with cells containing the WT LS and SS AGPase (Fig. S2). he LS^E370G^ mutation is shown on the 3D homology model of potato AGPase [34], where the residue appears not to be a part of the interface residues but rather is very close to the subunit interface residues (Fig. S1). We then decided to further characterize the function of LS-Glu370 in AGPase function.

### 3.2 Glutamate at position 370 in the LS modulates stability of heterotetrameric AGPase

We determined the heterotetrameric assembly properties of the LS^E370G^ AGPase with the wild type SS AGPase using native-PAGE and Western blot analysis. Before native-PAGE analysis, the integrity and expression levels of both subunits of LS^E370G^SS^WT^ and LS^WT^SS^WT^ AGPases were assessed by Western blot analyses. The LS and the SS proteins of the LS^E370G^SS^WT^ and wild type AGPases in cell free extract were detected around their predicted molecular mass (50 kDa), and their expression levels were comparable with the wild type AGPase (data not shown). Then, variable amounts (0.5, 1, and 2 μg of total protein) of the mutant and wild type AGPase samples were subjected to the 3-13% gradient native-PAGE followed by Western blot using anti-LS potato AGPase and anti-SS AGPase antibodies (Fig. 1A). The first uppermost bands correspond to approximately 200 kDa, which indicates the presence of tetramers. The second band indicates dimers, which were detected at approximately 100 kDa. The third band was identified at approximately 50 kDa and demonstrated the presence of monomers. On average, wild type AGPase heterotetramers comprised 30% of the total band intensity, whereas dimers and monomers comprised approximately 20% and 50% of the total band intensity, respectively, in both samples treated by anti-LS and anti-SS antibodies (Fig. 1B). A similar analysis of LS^E370G^SS^WT^ AGPase showed a drastic difference compared to the wild type AGPase. Almost 90% of the mutant AGPase was in the form of heterotetramers having comparable amount of the LS and the SS confirmed by anti-LS and anti-SS (Fig. 1B).

**Fig 1:**
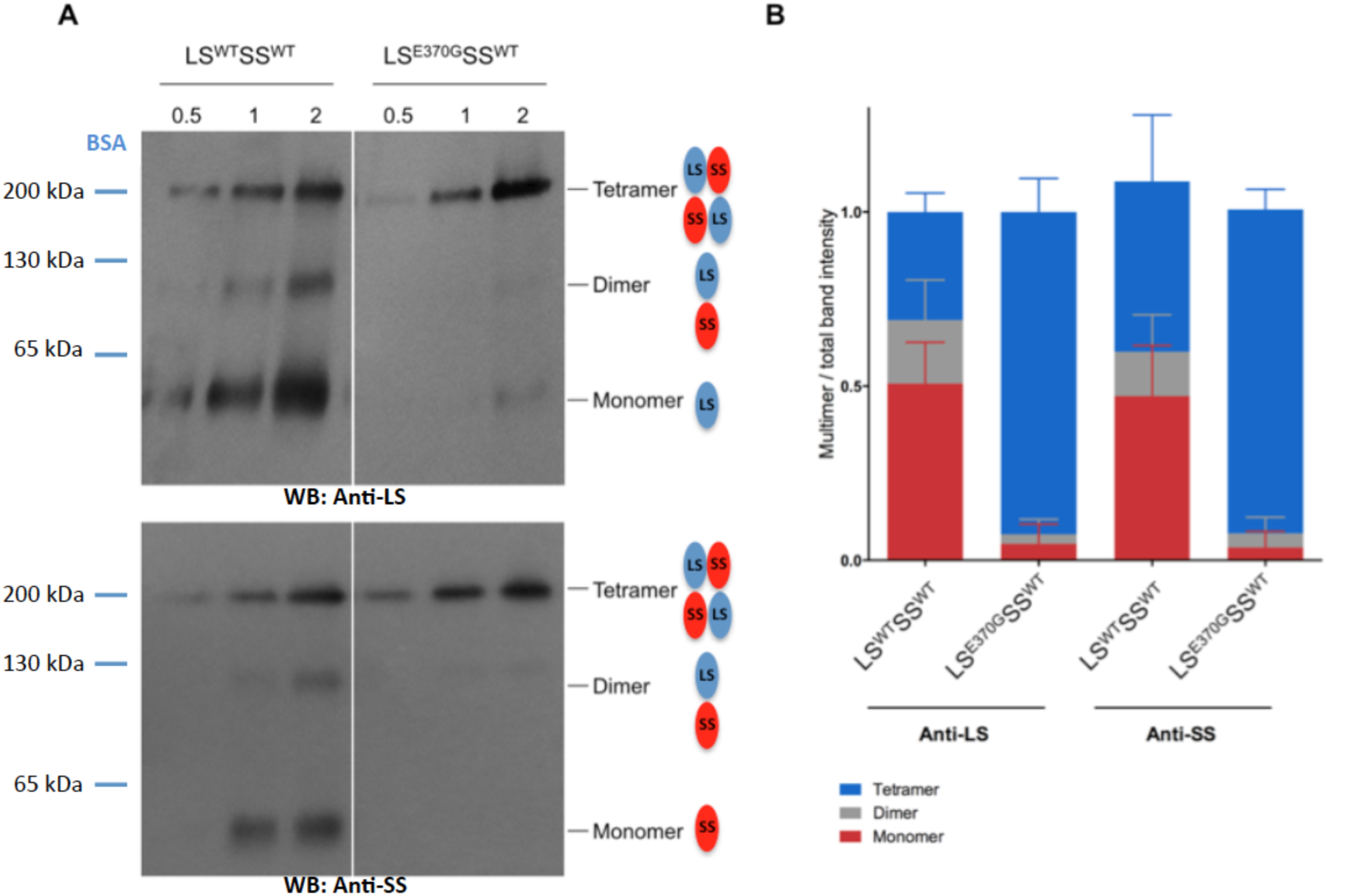
Native-PAGE followed by Western blot analysis of the LS^WT^SS^WT^ and LS^E370G^ SS^WT^ AGPases. (A) Crude extract samples were loaded in equal increasing amounts 0.5, 1 and 2 μg of total protein. 3-13% gradient native polyacrylamide gels were used to separate monomeric, dimeric and tetrameric forms of AGPase. Western blot was performed using either potato specific LS and SS antibodies. (B) A stacked column graph showing the ratio of monomer, dimer and tetramer to total band intensity, respectively. Average values with standard error for >4 independent experiments were compared with Student’s t test with regard to the wild type. ***P <0.0001; *P<0.05.

These results indicated that comparable amount of heterotetrameric AGPase were formed in both wild type and mutant AGPases. However, the amount of the dimer and monomer of the LS along with the SS was dramatically reduced compared to wild type AGPase. To see the how Glu at the position of 370 might affect the disappearance of the LS in monomer and dimer bands we performed charge distribution analysis on the region where LS and SS interacts in head and tail orientation on 3D-modeled heterotetrameric potato AGPase (Fig. 2A). The presence of the Gly instead of the Glu at the position of 370 in LS drastically changes surface charge distribution towards positive around that region (Fig. 2B). Then, we decided to replace Glu-370 with Arg in the LS AGPase to further shift surface towards more positive and investigate the effect of such mutation on the heterotetrameric stability of the AGPase. Native-PAGE analysis revealed that LS^E370R^SS^WT^ AGPase has enhanced LS and SS dimer formation rather than heterotetrameric assembly (Fig. 3A). The quantity of the gel showed that the second band, indicating the presence of dimers, consists of 40%, whereas the amount of the dimer in wild type AGPase was approximately 20% (Fig. 3B). All these experiments let us hypothesize that a change in the surface charge of LS might affect the stability of the AGPase.

**Fig 2:**
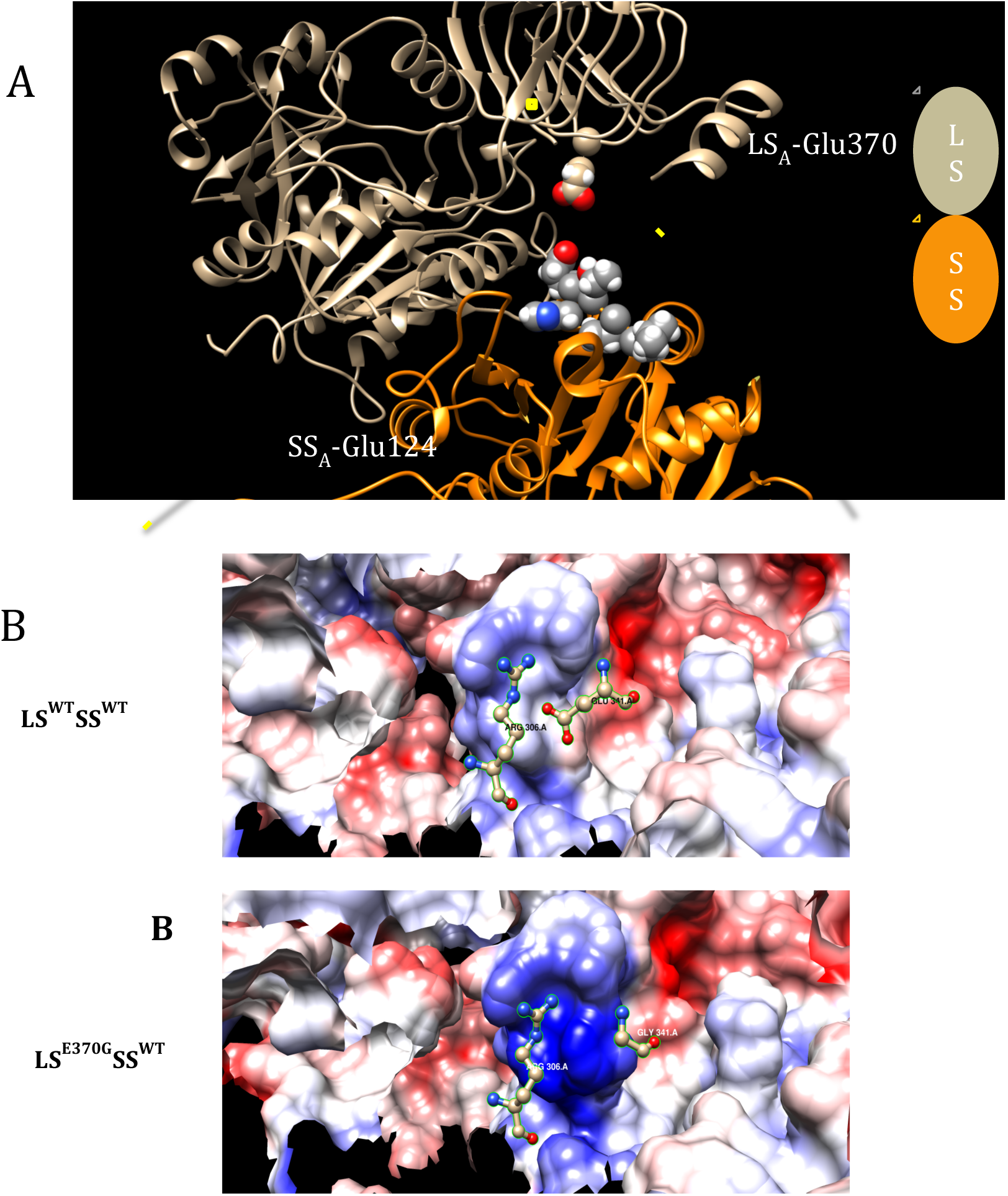
Surface charge analysis of the interaction site in the 3D modeled (A) LS^WT^SS^WT^ and (B) LS^E370G^SS^WT^ AGPases.

**Fig 3:**
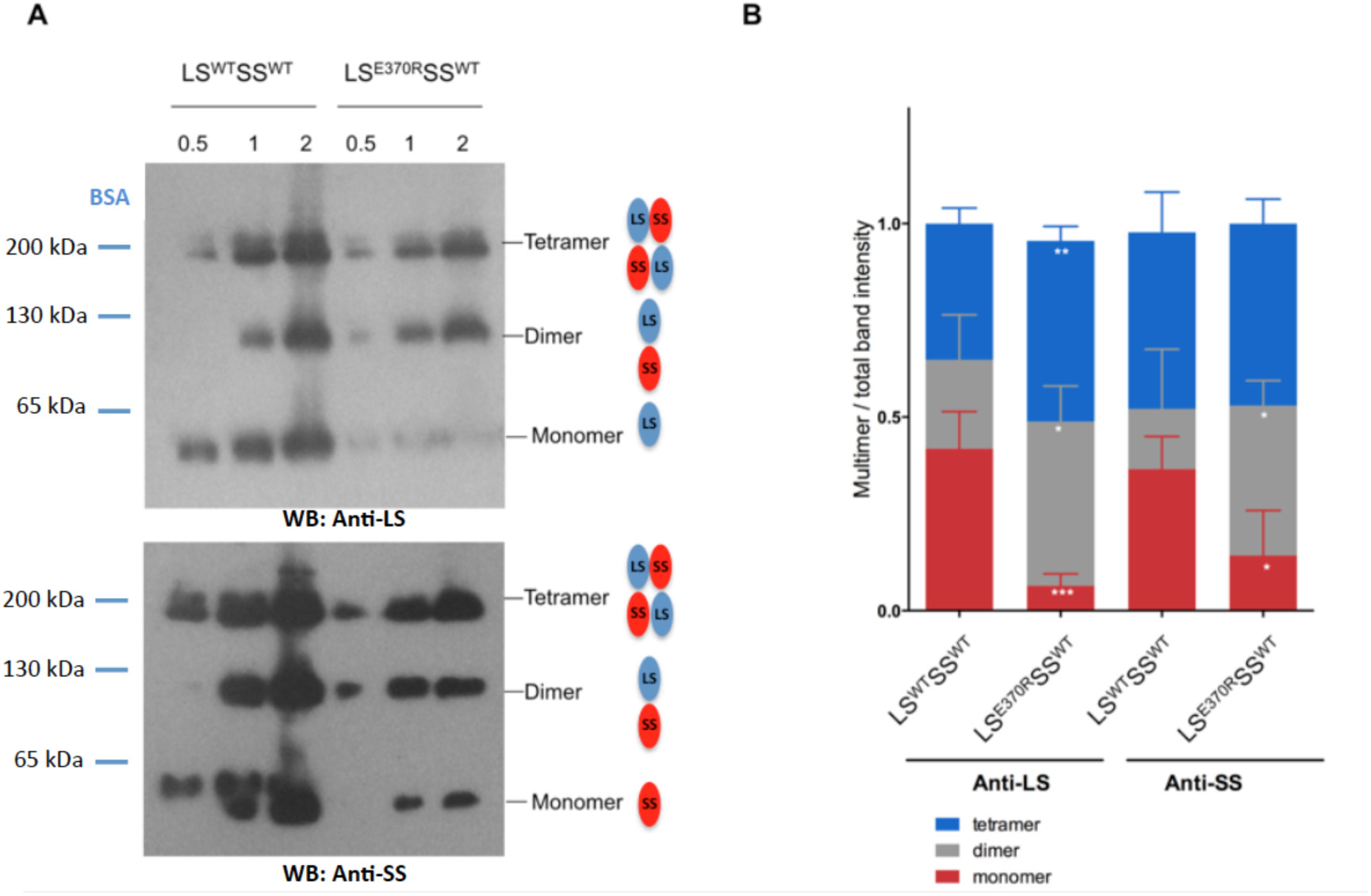
Native-PAGE followed by Western blot analysis of the LS^WT^ SS^WT^ and LS^E370R^ SS^WT^ AGPases. (A) Crude extract samples were loaded in equal increasing amounts 0.5, 1 and 2 μg of total protein. Western blot was performed using either potato specific LS and SS antibodies. (B) A stacked column graph showing the ratio of monomer, dimer and tetramer to total band intensity, respectively. Average values with standard error for >4 independent experiments were compared with Student’s t test with regard to the wild type. ***P <0.0001; *P<0.05.

Computational analysis was performed to understand how the LS^E370G^ and LS^E370R^ mutations affect stability of the LS AGPase. Over the last decade various programs were developed to predict the protein stability upon point mutations by applying different approaches, namely EGAD [49], I-Mutant [50], CC/PBSA [51], FoldX [52], Rosetta [53], SDM [53], POPMUSIC [54] and others. FoldX produces quantitative estimation of protein stability upon mutation [43] and calculates the interaction energies between the subunits of a protein complex accurately [44–46]. We therefore used FoldX to calculate the effect of point mutations on the stability of LS^E370G^ and LS^E370R^ (ΔΔG), and also on the overall interaction energy in heterotetrameric AGPase. FoldX analysis indicated that mutation of LS^E370G^ (ΔΔG= 4.1kcal/mol) and LS^E370R^ (ΔΔG=5.1 kcal/mol) causes destabilization of the subunit (ΔΔG > 0 shows destabilizing effect and ΔΔG < 0 shows stabilizing effect (Equation 1)). This also indicates that as the side chain positivity of 370^th^ residue increases, the stability of the LS decreases.

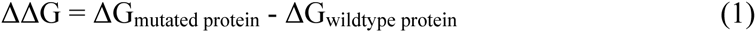

A detailed analysis of the energy decomposition of ΔΔG indicated that followings are the main factors contributing instability of the mutant LS AGPases: Side-chain Hbond, van der Waals, electrostatics and solvation hydrophobic terms contribute as the destabilizing factors (for each factor ΔΔG > 1kcal/mol) in LS^E370G^ (Table S2). These results imply that hydrogen-bond network and electrostatic interactions around the 370^th^ residue are disturbed in both mutants causeing less stable LS AGPase. To further validate these results, we used I-Mutant 3.0 (structure mode) [47] and POPMUSIC server (Table S3) where they consistently show that these mutations have destabilizing effect on the LS. All these computational results explain how 370^th^ residue contributes the stability of the LS AGPase.

Then we would like to calculate the interaction energies between subunits of AGPase in wildtype and mutants to investigate the effect of mutations on the interaction of subunits. Our results (Table 1) show that LS^E370G^ mutation does not cause overall interaction energy change between the subunits of AGPase (Table S4). On other hand LS^E370R^ mutation slightly decreases the interaction energy and indicates that mutation favors the formation of the heterotetrameric AGPase (Table S5).

**Table 1:** ΔΔG values of interaction energies between subunits of AGPase produced by FoldX. ΔTm value of mutant AGPases produced by HOTMUSIC.

| | $\Delta\Delta G$ (kcal/mol) | $\Delta T_m$ (K) |
| --- | --- | --- |
| LS<sup>E370G</sup>AGPase | $0.0 \pm 0.04$ | -4.96 |
| LS<sup>E370R</sup>AGPase | $-0.7 \pm 0.1$ | -0.52 |

According to the heterotetrameric AGPase model [30, 32–34], such interactions are only possible between LS and SS in a head to tail orientation (Fig. 2). On the other hand, mutant LS^R370^, may interact with SS^E124^ because of the opposite charged amino acids on these subunits and this may elevate their affinities to each other and result more LS-SS dimer formation. Our computational analysis also indicated that this LS^R370^ mutation favors head to tail interaction between LS^R370^ and the SS (Table S5). This might explain why we observed more dimer formation between LS^R370^ and the SS (Fig. 2A).

### 3.3 Heterotetramers of LS^E370G^ SS^WT^ AGPase are heat labile

We used a tag-less expression system described in [13] to investigate the effect of temperature on the enzyme stability and activity. Cell free extracts of the wild type and mutant AGPases were incubated at 65°C for 5 and 10 minutes. Then, samples were centrifuged and subjected to the 3-13% native-PAGE followed by Western blot. Fig. 4 clearly showed that the heterotetrameric form of the wild type AGPase was preserved while the heterotetrameric form of the LS^E370G^ SS^WT^ AGPase totally disappeared at the end of the heat treatment. We also tested the heat stability of LS^E370R^ SS^WT^ AGPase under the same conditions. The heterotetrameric assembly of the mutant AGPase disappeared at the end of heat treatment, which was similar to the LS^E370G^ SS^WT^ AGPase (Fig. S3). hen, the samples were used to measure the remaining activities at saturated concentrations of substrates (5mM) and 3PGA (20mM) with forward direction assay. There were no detectable remaining activities of LS^E370G^ SS^WT^ and LS^E370R^ SS^WT^ (data not shown).

**Fig 4:**
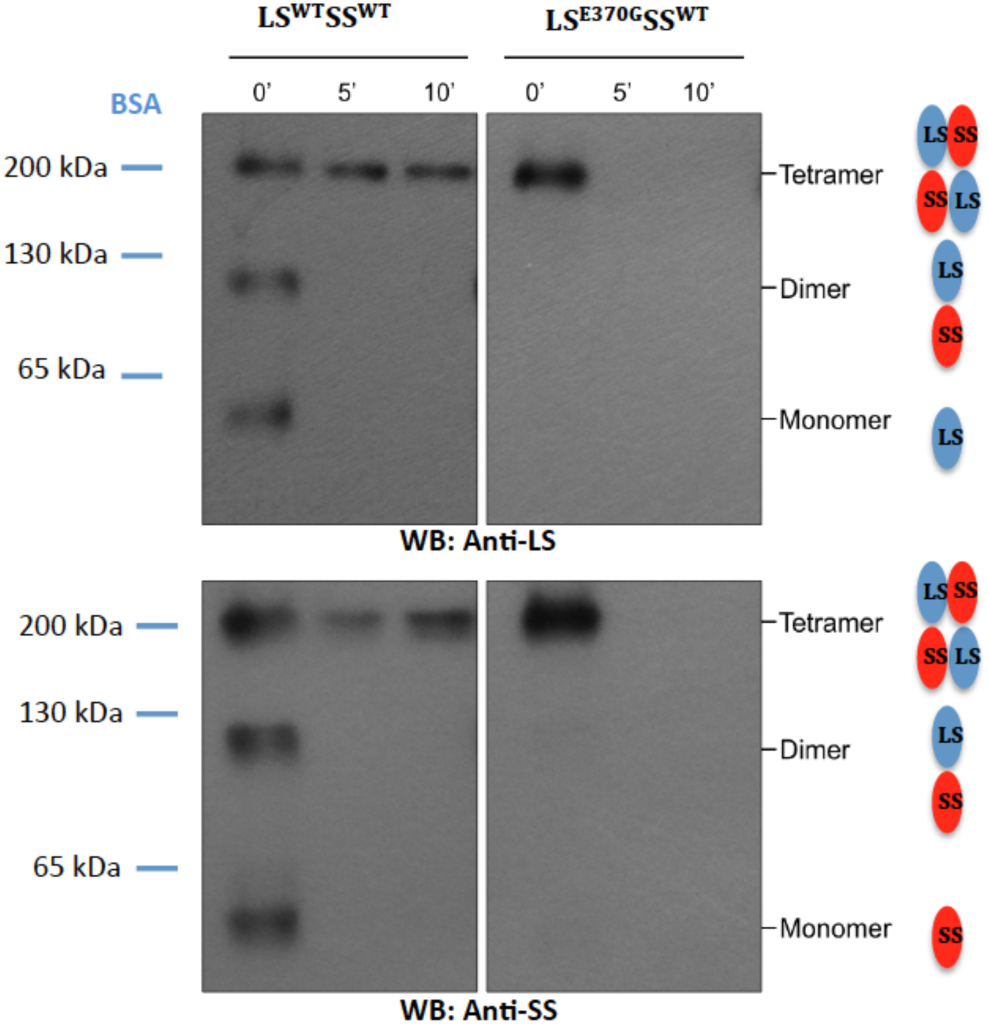
Comparing heat stability of LS^E370G^SS^WT^ with the wild type AGPase at 65°C. (Native PAGE analysis of heat-treated samples for 5 and 10 min at 65°C. The same amount and same volume of total soluble protein samples were loaded into 3-13% native polyacrylamide gels and western blots were detected using either potato specific LS and SS antibodies.

To explore the possibility that the LS^E370G^SS^WT^ AGPase has altered redox status mediated by Cys-12 it was subjected to non-reducing SDS–PAGE followed by Western blot. The results indicated that both WT and LS^E370G^SS^WT^ AGPases had all LS subunits as monomers and had comparable disulfide bond between the SS (Fig. S4).

Finally we investigated the heat stability of our mutant AGPases by using the recently developed HOTMUSIC tool which predicts melting temperature changes as a result of point mutations [6]. HOTMUSIC analysis indicated that both mutations cause protein to adopt lower Tm which means decrease in the heat stability (Table 1).

### 3.4 Kinetic and allosteric characterization

Polyhistidine tagged heterotetrameric AGPases were purified using DEAE weak anion exchange chromatography followed by TALON metal affinity chromatography with a final purity of 60-70%. The purified recombinant wild type and mutant AGPases were assayed in the forward reaction direction to determine the effects of mutations on the enzyme activity. The kinetic and allosteric parameters of the wild type and mutant AGPases are shown in the Table 2. Kinetic properties of wild type AGPase were comparable with previously published kinetic values of the recombinant heterotetrameric AGPase [13].

Analysis of the substrate binding properties of the AGPases showed that there were no significant differences in the ATP K_m_ values between the heterotetrameric wild type and LS^E370G^SS^WT^ AGPases (Fig. S5A). In contrast, the affinity of the enzymes towards the other substrate G1P showed more variation. The wild type AGPase has a K_m_ for G1P of 0.044 mM. However, the G1P affinity of LS^E370G^SS^WT^ AGPase was approximately 3-fold lower than of the wild type, with a K_m_ of 0.13 mM (Fig. S5B).

**Table 2:**
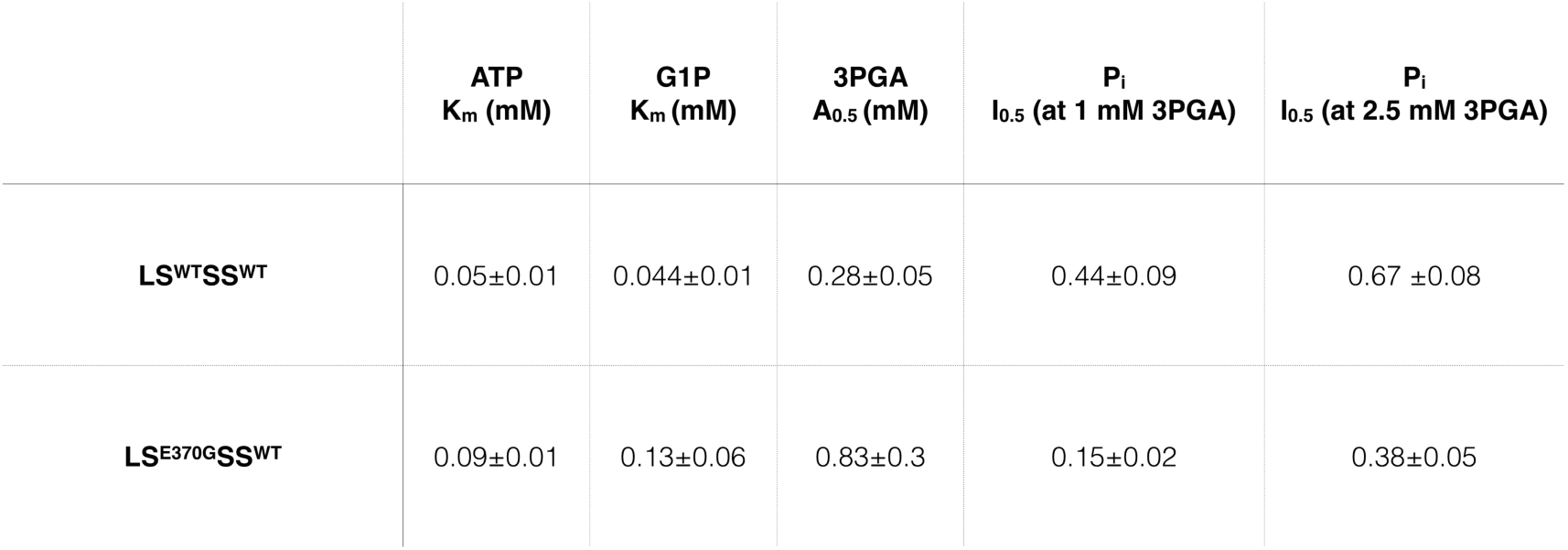
Kinetic and allosteric parameters of the wild type and LS^E370G^SS^WT^ AGPase. Kinetic and regulatory properties were determined in the ADP-glucose synthesis direction. All values are in mM. K_m_ and A_0.5_ for substrates and 3PGA, respectively, correspond to the concentration of these molecules required for the enzyme activity to attain 50% of maximal activity. I_0.5_ is the amount of P_i_ required to inhibit the enzyme activity by 50% of maximal activity in the presence of either 1 mM or 2.5 mM 3PGA. Depicted are the mean values with standard error of at least two independent experiments.

Allosteric regulatory properties varied considerably between the wild type and mutant AGPases. The A_0.5_ values for 3PGA with wild type and LS^E370G^SS^WT^ AGPases are 0.28 and 0.83 mM, respectively (Fig. S6A). At a lower 3PGA concentration (1 mM), the LS^E370G^ SS^WT^ mutant showed increased sensitivity towards Pi inhibition compared with the wild type AGPase. The I0.5 of the LS^WT^ SS^WT^ AGPase was 0.44 mM; meanwhile, it was 0.15 mM for the LS^E370G^ SS^WT^ mutant. The LS^E370G^ SS^WT^ AGPase was still more sensitive to the Pi inhibition compared with the wild type enzyme with I0.5 values 0.38 and 0.67 mM, respectively, in the presence of 2.5 mM 3PGA (Fig. S6B,C).

Our results indicated that the structural change due to the E370G mutation in the LS affects the kinetic and allosteric properties of the heterotetrameric AGPase. Previous studies have shown the importance of the LS region close to the subunit interface residues for the allosteric regulation of AGPase [27, 33, 41, 55, 56]. These data are in agreement with the previous studies indicating that both the LS and SS are equally important in enzyme catalysis and regulation [55–57].

To determine the degree of the conservation of the Glu residue at position 370 in the LS, multiple alignments from the mature LS sequences of various plants were performed using CLUSTAL (Fig. S7). Conservation analysis revealed that the glutamate residue at position 370 in the LS is not highly conserved among different plant species. However, the neighboring amino acids are conserved, and valine and isoleucine residues are mostly observed at this position. In fact the maize counterpart of this residue was previously identified with a phylogenetic analysis as a candidate residue (positively selected type II site), which might have an important effect for the function of the protein [33, 57].

Maize LS^V4161^ AGPase leads to increased heat stability without affecting the kinetic constants, whereas maize LS^V4161^ AGPase significantly impairs 3PGA binding but does not change the heat stability. Those together with our results, which reveal the contribution of Glu370 for the heat stability, substrate and effector binding properties, specify the importance of this residue for the structure-function relationship of AGPase. Additionally this study showed that Glu370 affects oligomeric status of the AGPase by enhancing the formation of the LS-SS dimer. In 3D homology modeled potato heterotetrameric AGPase we observed that replacement of Glu into Gly drastically shifted surface charge distribution towards positive in the LS and this region is close proximity with SS^E124^ with distance 5Å (Fig. 2A). It is most likely that SS^E124^ strongly interacts with this positively charged region and dimer formation between the LS^E370R^ and WT SS yet to be shown by crystal structure study. Further kinetic characterization showed that the LS^E370G^ SS^WT^ AGPase has reduced affinity in both substrate (G1P) and activator (3PGA). Therefore, the expression of the LS^E370G^ AGPase with SS^WT^ AGPase in bacterial system fails complement glgC in *E.coli* (Fig S2).

In conclusion, our results show that LS^E370^ intricately modulate the heterotetrameric assembly, heat stability and enzymatic activity of the AGPase.

## 4. Acknowledgment

This work was supported by TUBITAK-KBAG 114Z760; TUBITAK-BIDEP 2211 [PhD fellowship of A.B.S].

---

**Figure S1:**
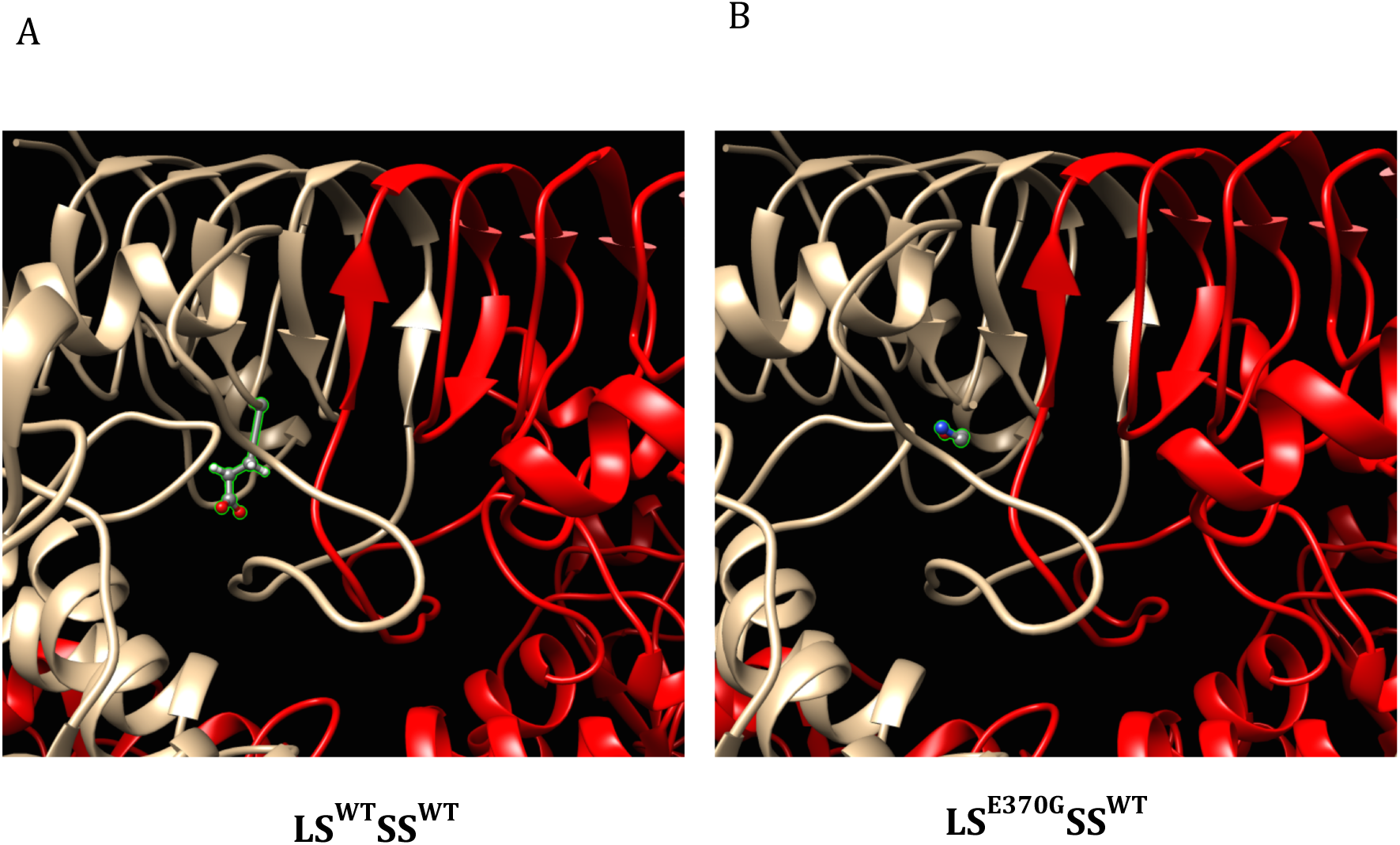
Location of the wild type and mutant (E370G) residues at position 370 in the LS on homology modeled AGPase in which the LS and the SS are depicted by tan and red colors, respectively. LS^370^ residue is colored according to its elements

**Figure S2:**
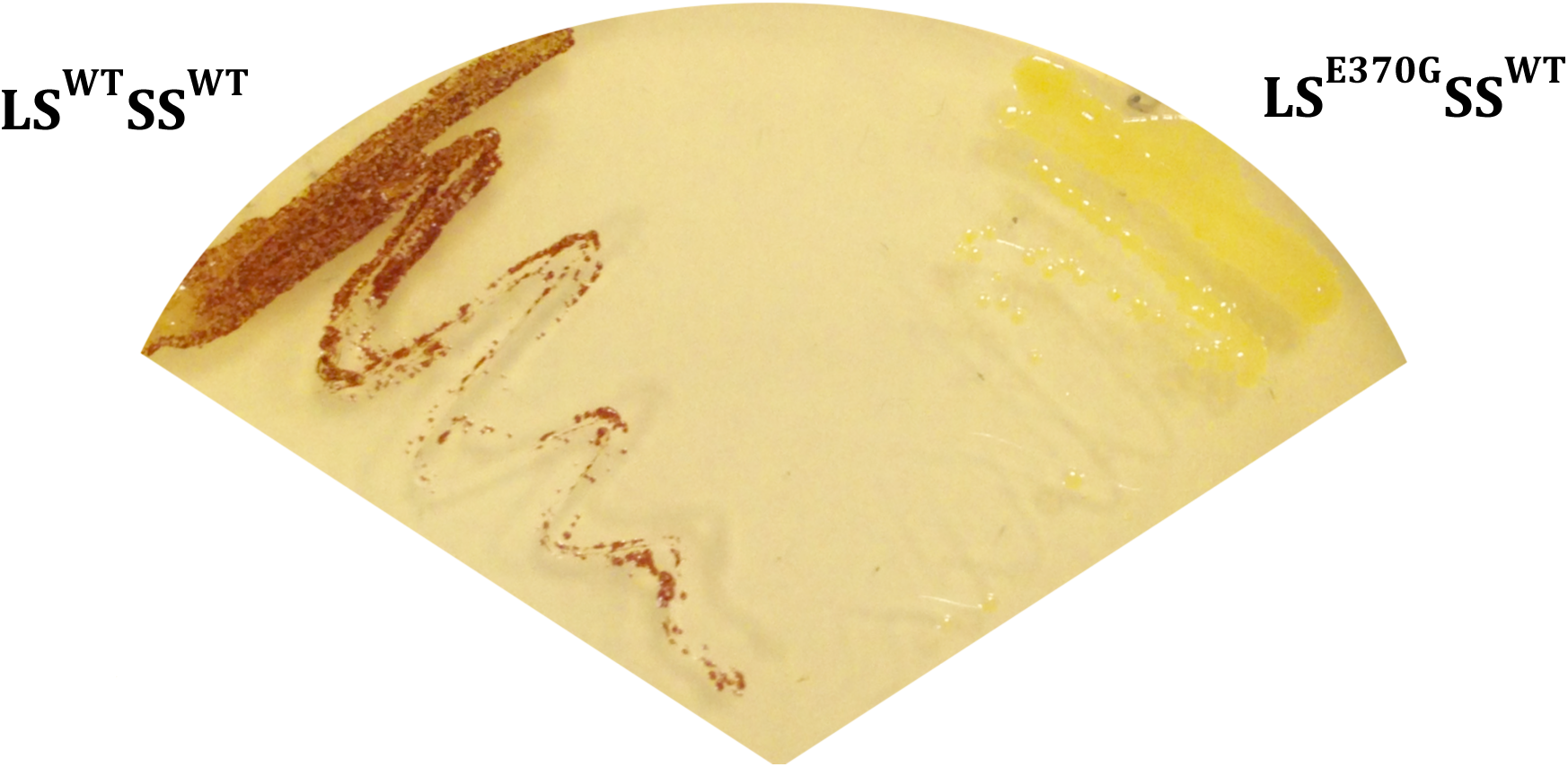
Bacterial complementation assay by iodine staining of *E.coli* glgC-cells expressing the potato tuber recombinant LS^WT^SS^WT^ and LSE^370G^SS^WT^ AGPases. The plate was streaked from a single colony of each strain onto Kornberg’s 1% glucose enriched plate and incubated overnight at 37°C

**Figure S3:**
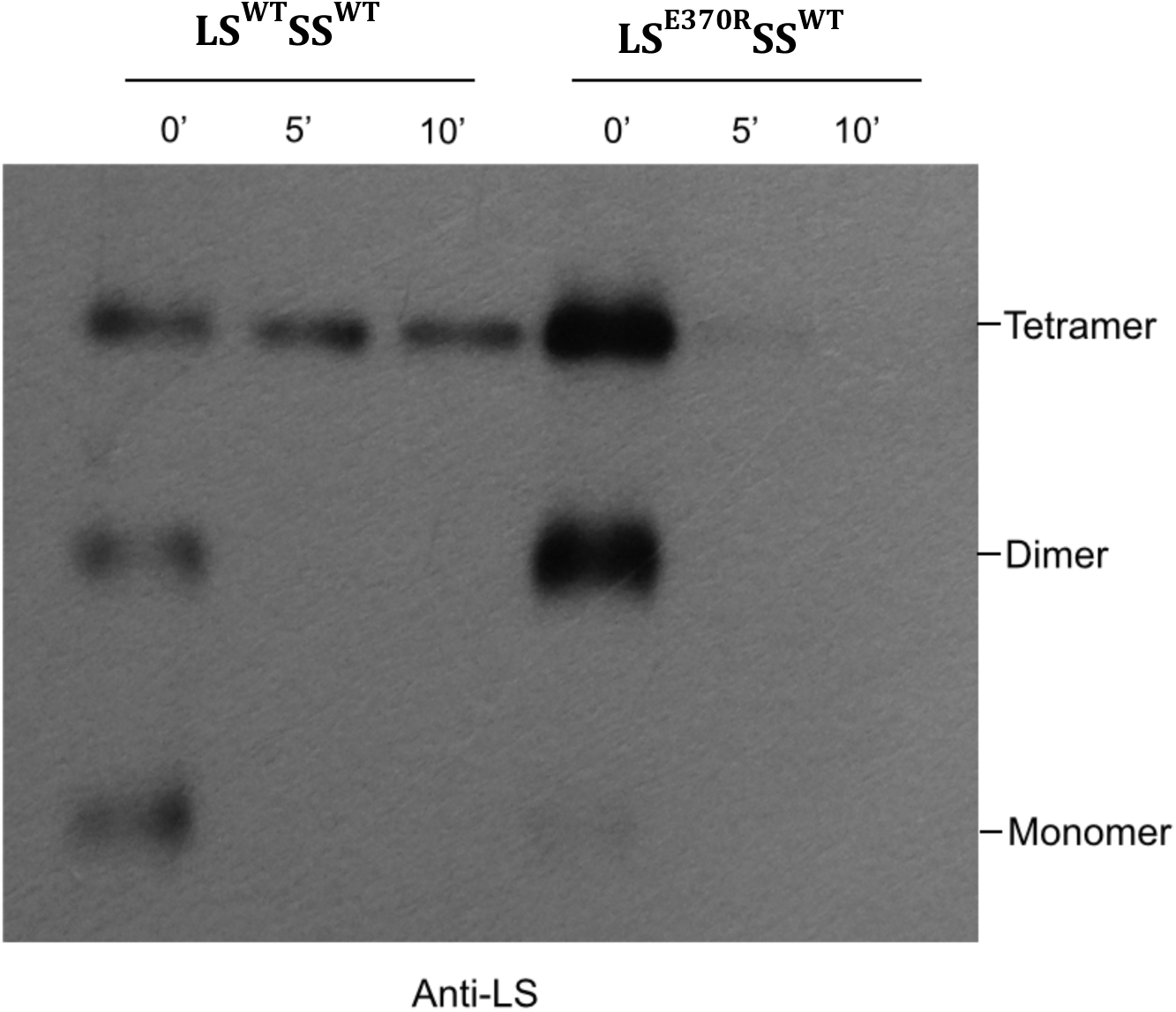
Comparing heat stability of LS^E370R^SS^WT^ with the wild type AGPase at 65°C. Native PAGE analysis of heat-treated samples for 5 and 10 min at 65°C. The same amount and same volume of total soluble protein samples were loaded into 3-13% native polyacrylamide gels and western blots were detected using potato specific LS antibody.

**Figure S4:**
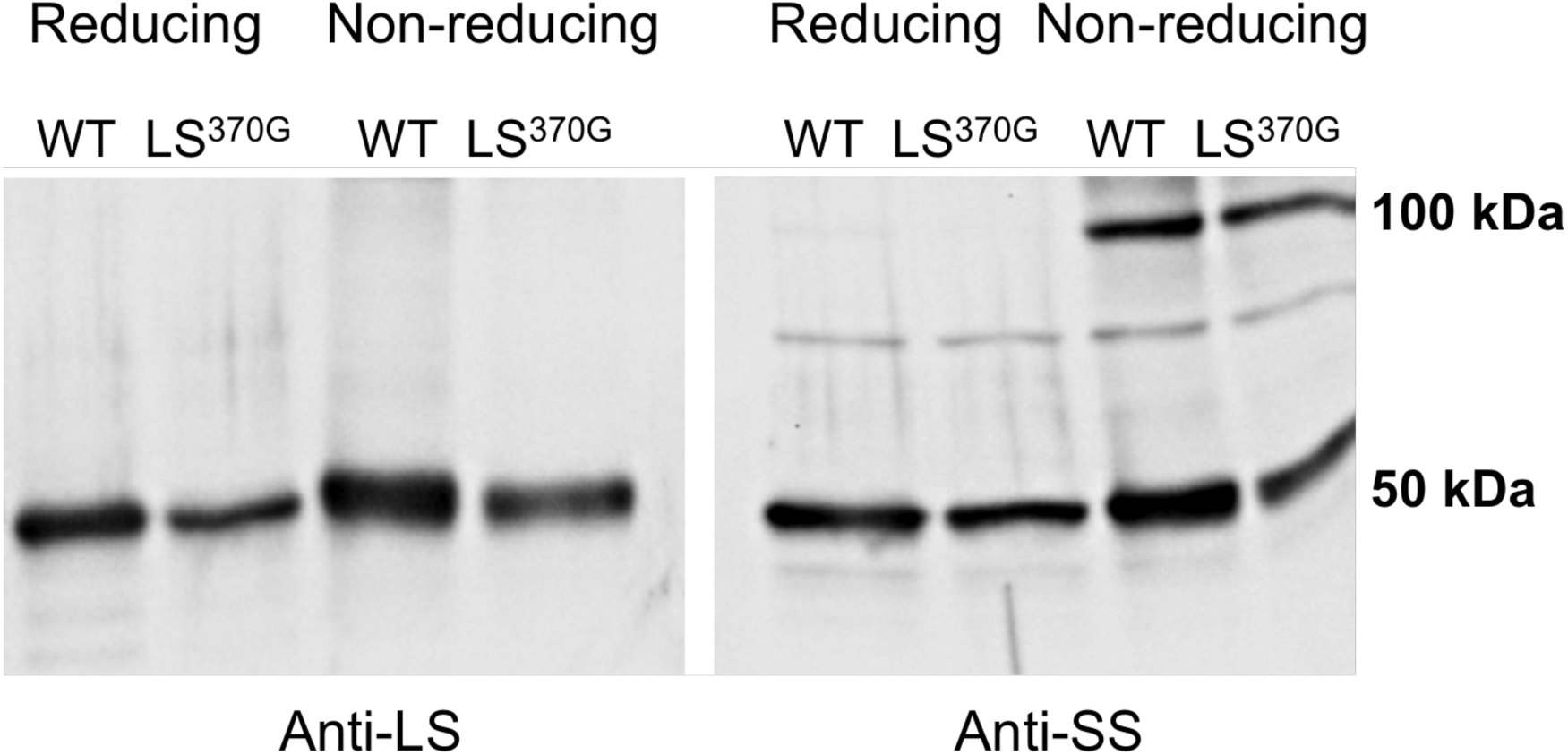
Comparison of reducing and non-reducing SDS-PAGE western blot analysis of the wild-type and E370G mutant. A non-reducing 10% SDS-PAGE gel followed by western blot with potato specific anti-LS and anti-SS antibodies was performed to investigate the formation of the disulfide bridge. Only anti-SS probed blot showed dimers in both the wild type and LS^E370G^ AGPase.

**Figure S5:**
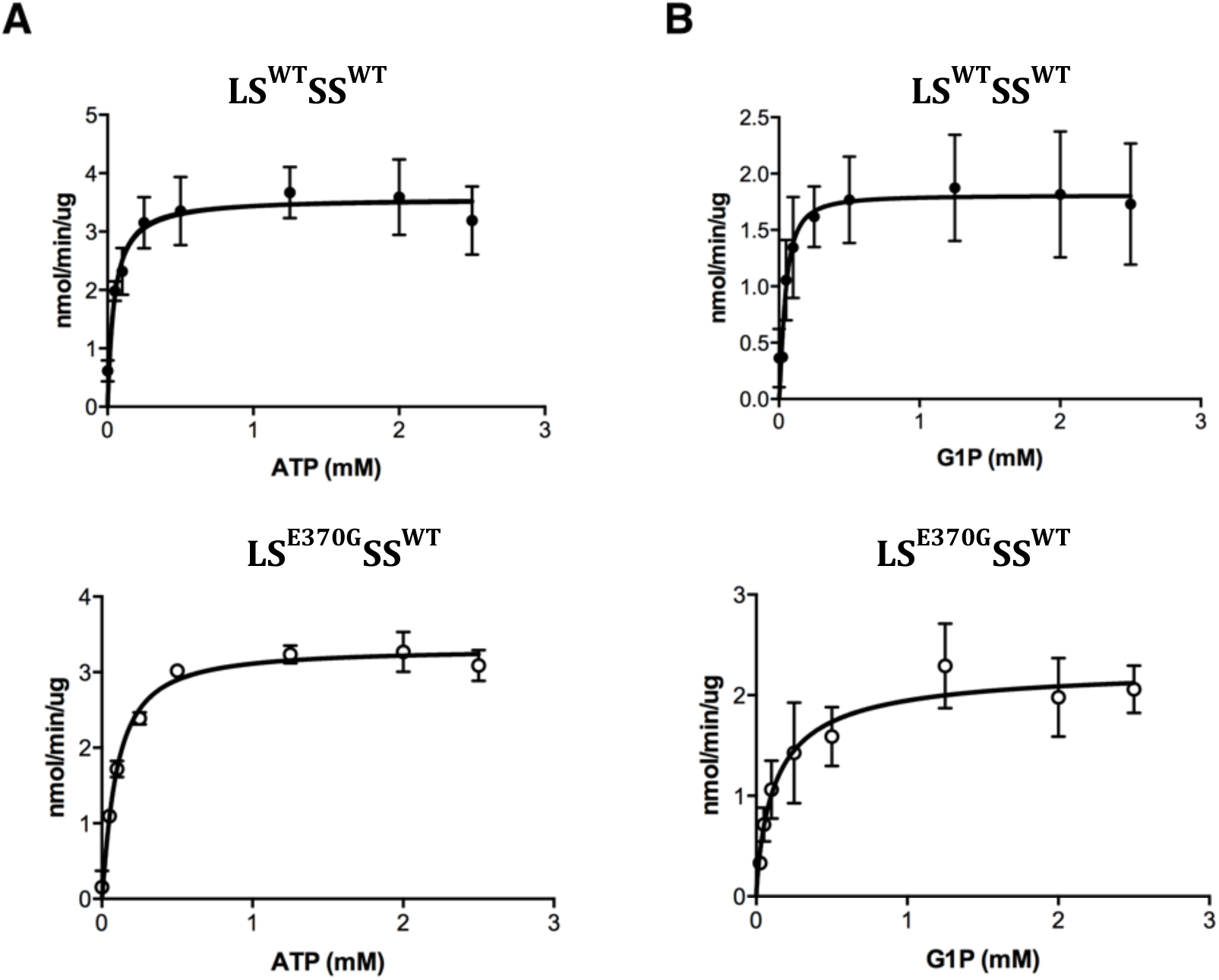
(A) Velocity versus ATP concentration graphs of the wild type and LS^E370G^SS^WT^ AGPases in the presence of 5 mM 3PGA. (B) Velocity versus G1P concentration graphs of the wild type and LS^E370G^ SS^WT^ AGPases in the presence of 5 mM 3PGA.

**Figure S6:**
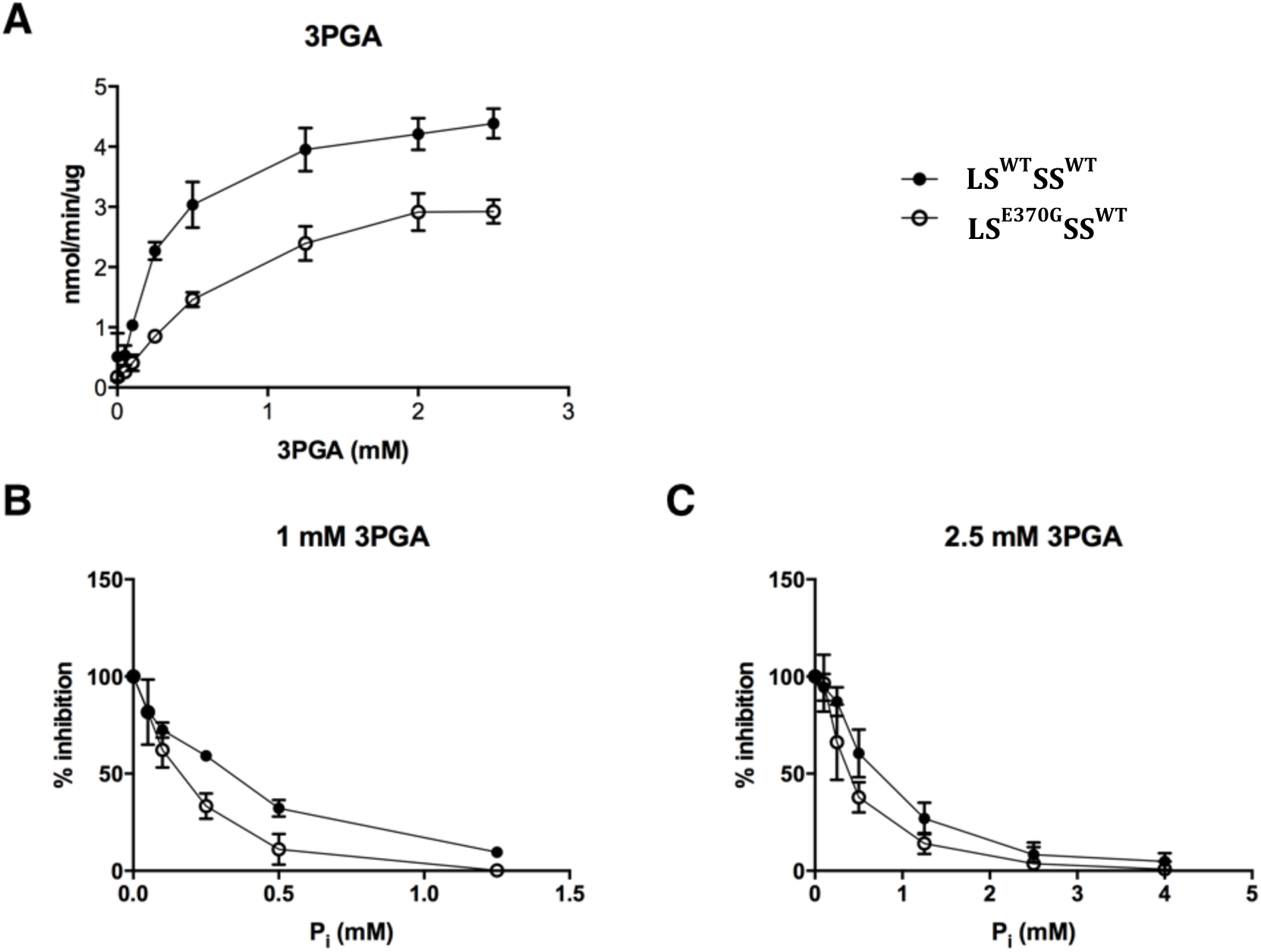
3PGA activation (A) and P_i_ inhibiton (B,C) profiles of the wild type (•)AGPase and LS^E370G^SS^WT^ (¢)mutant AGPase. (A) Reactions were performed in the forward direction under saturating substrate conditions and with varying amounts of 3PGA (B) Reactions were performed in the forward direction under saturating substrate conditions and with 1 mM 3PGA. (C) Reactions were performed in the forward direction under saturating substrate conditions and with 2.5 mM 3PGA.

**Figure S7:**
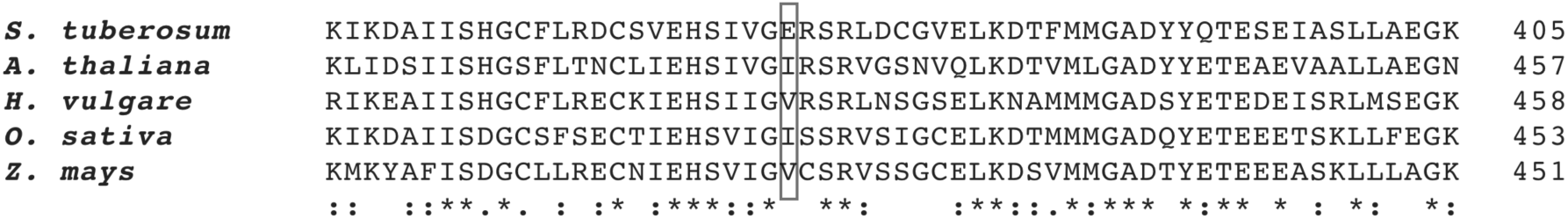
Primary sequence alignments of AGPase LSs from different plant species. Box indicates the residue investigated in this study. *S. tuberosum* (Potato LS; Q00081), *A. thaliana* (Arabidopsis; AAB58475), *H. vulgare* (Barley; CAA47626), *O. sativa* (Rice; BAG92523), *Z. mays* (Maize; P55241)

**Table S1:**
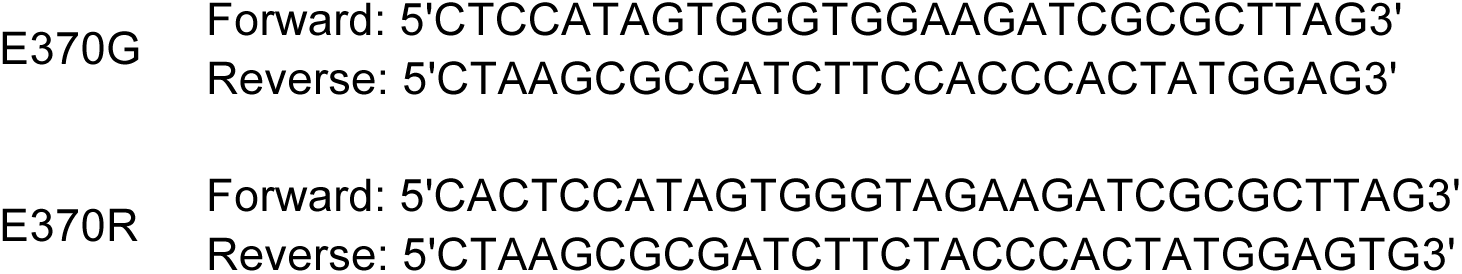
Primer sequences used in this study for site-directed mutagenesis

**Table S2:**
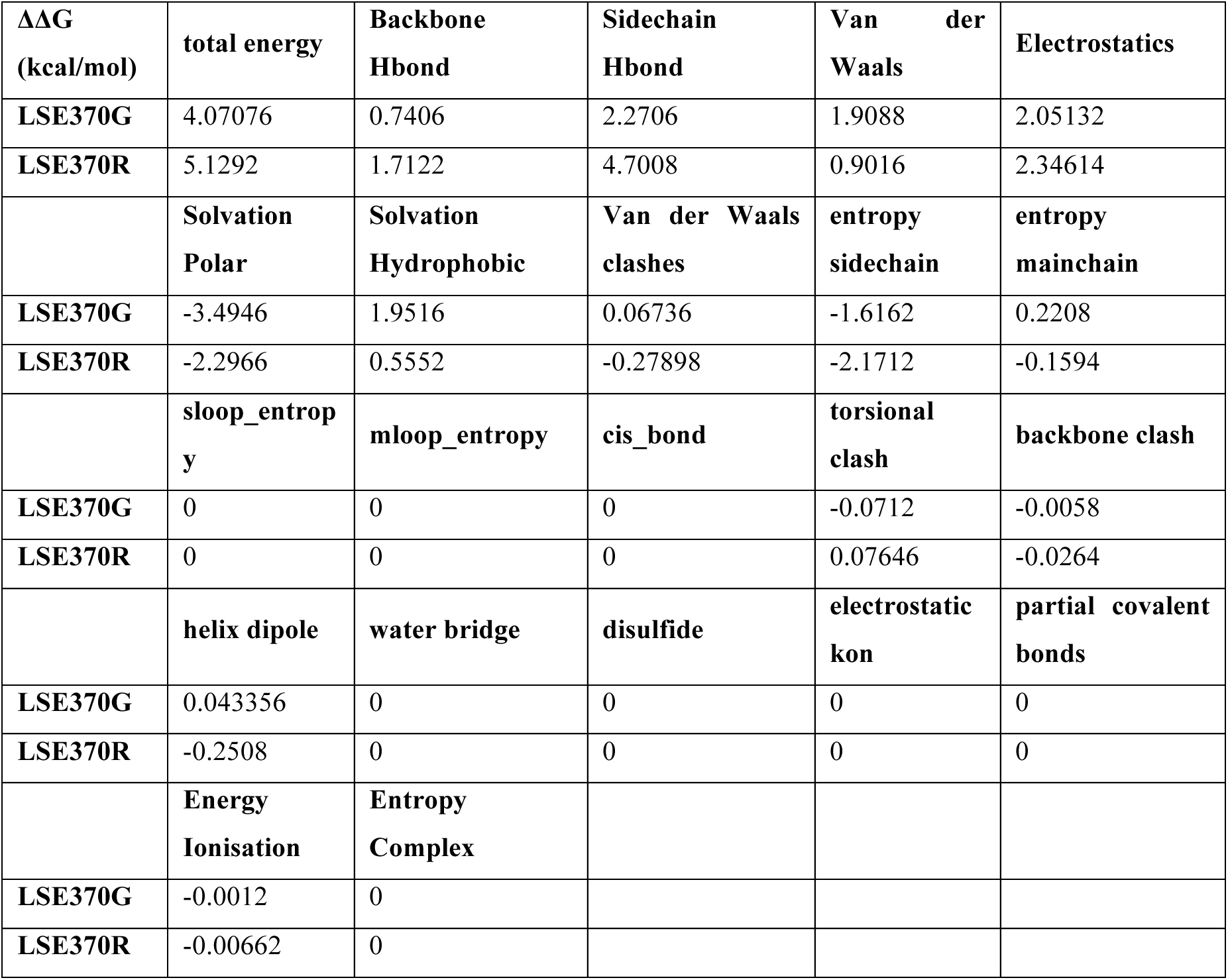
Energy decomposition of ΔΔG values of mutant LSs.

| **$\Delta\Delta G$**<br>**(kcal/mol)** | **total energy** | **Backbone**<br>**Hbond** | **Sidechain**<br>**Hbond** | **Van der**<br>**Waals** | **Electrostatics** |
| --- | --- | --- | --- | --- | --- |
| **LSE370G** | 4.07076 | 0.7406 | 2.2706 | 1.9088 | 2.05132 |
| **LSE370R** | 5.1292 | 1.7122 | 4.7008 | 0.9016 | 2.34614 |
|  | **Solvation**<br>**Polar** | **Solvation**<br>**Hydrophobic** | **Van der Waals**<br>**clashes** | **entropy**<br>**sidechain** | **entropy**<br>**mainchain** |
| **LSE370G** | -3.4946 | 1.9516 | 0.06736 | -1.6162 | 0.2208 |
| **LSE370R** | -2.2966 | 0.5552 | -0.27898 | -2.1712 | -0.1594 |
|  | **sloop_entropy** | **mloop_entropy** | **cis_bond** | **torsional**<br>**clash** | **backbone clash** |
| **LSE370G** | 0 | 0 | 0 | -0.0712 | -0.0058 |
| **LSE370R** | 0 | 0 | 0 | 0.07646 | -0.0264 |
|  | **helix dipole** | **water bridge** | **disulfide** | **electrostatic**<br>**kon** | **partial covalent**<br>**bonds** |
| **LSE370G** | 0.043356 | 0 | 0 | 0 | 0 |
| **LSE370R** | -0.2508 | 0 | 0 | 0 | 0 |
|  | **Energy**<br>**Ionisation** | **Entropy**<br>**Complex** |  |  |  |
| **LSE370G** | -0.0012 | 0 |  |  |  |
| **LSE370R** | -0.00662 | 0 |  |  |  |

**Table S3:** ΔΔG values of two different individual mutants of LS produced by POPMUSIC and I-Mutant 3.0. I-Mutant 3.0 calculations classify the results as large decrease in stability if ΔΔG > 0.5kcal/mole.

| | $\Delta\Delta G$ (kcal/mol) | $\Delta\Delta G$ (kcal/mol) |
| --- | --- | --- |
|  | From POPMUSIC | From I-Mutant 3.0 |
| LS<sup>E370G</sup> | 1.75 | 2.03 |
| LS<sup>E370R</sup> | 0.44 | 1.05 |

**Table S4:**
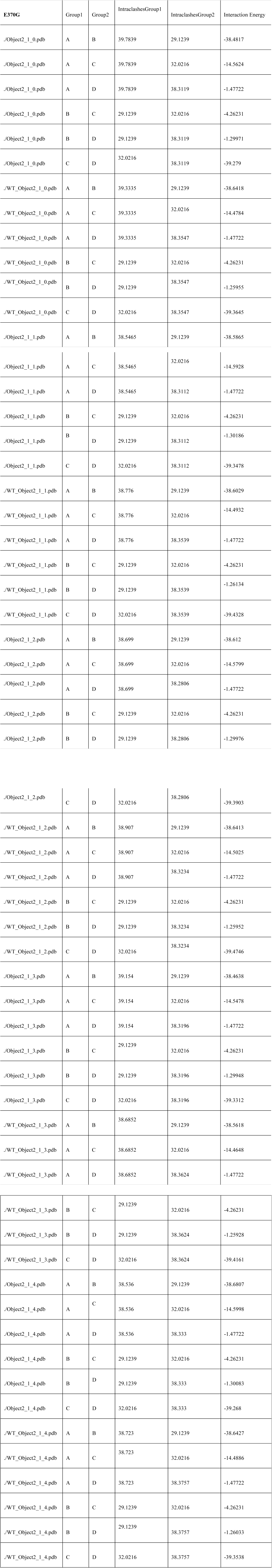
Interaction energies of subunits of AGPase in LSE370G and wildtype AGPase. Object2 represents the LSE370G and WT_Object represents the wildtype AGPase. The same calculation was run 5times to make sure algorithm converged. A and D are LSs; B and C are SSs

**Table S5:**
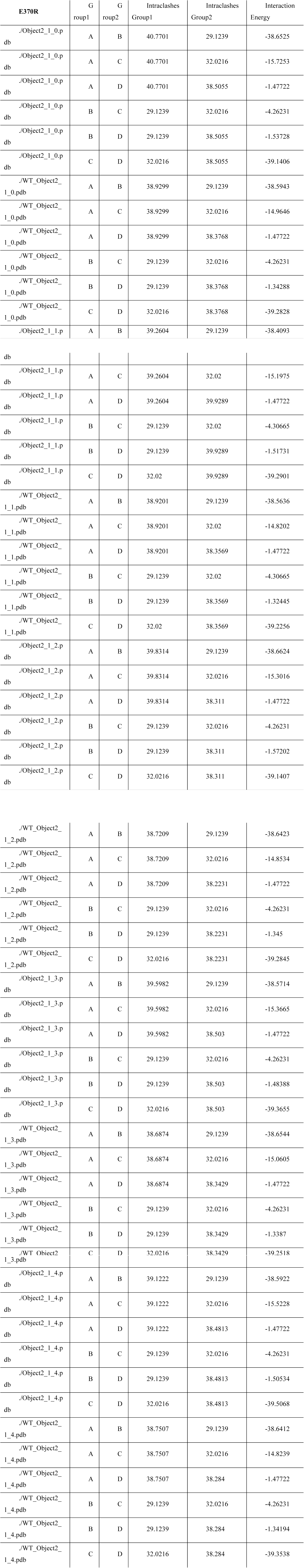
Interaction energies of subunits of AGPase in LSE370R and wildtype AGPase. Object2 represents the LSE370R and WT_Object represents the wildtype AGPase. The same calculation was run 5times to make sure algorithm converged. A and D are LSs; B and C are SSs. Head and tail interaction between the LS and the SS indicated by A and C, respectively To observe the effect of mutation on dimer formation, compare the interaction energy results of A and C in Object2_1_0.pdb and WT_Object2_1_0.pdb.

## References

[1] C.J. Slattery, I.H. Kavakli, T.W. Okita, Engineering starch for increased quantity and quality, Trends Plant Sci, 5 (2000) 291–298.

[2] M.A. Ballicora, A.A. Iglesias, J. Preiss, ADP-glucose pyrophosphorylase: a regulatory enzyme for plant starch synthesis, Photosynth Res, 79 (2004) 1–24.

[3] I.H. Kavakli, C.J. Slattery, H. Ito, T.W. Okita, The conversion of carbon and nitrogen into starch and storage proteins in developing storage organs: an overview, AustJ Plant Physiol, 27 (2000) 561–570.

[4] J.R. Sowokinos, Pyrophosphorylases in Solanum-Tuberosum .2. Catalytic Properties and Regulation of Adp-Glucose and Udp-Glucose Pyrophosphorylase Activities in Potatoes, Plant Physiol, 68 (1981) 924–929.

[5] J.R. Sowokinos, J. Preiss, Pyrophosphorylases in Solanum-Tuberosum .3. Purification, Physical, and Catalytic Properties of Adp-Glucose Pyrophosphorylase in Potatoes, Plant Physiol, 69 (1982) 1459–1466.

[6] A. Tuncel, T.W. Okita, Improving starch yield in cereals by over-expression of ADPglucose pyrophosphorylase: expectations and unanticipated outcomes, Plant Sci, 211(2013)52–60.

[7] M.R. Bhave, S. Lawrence, C. Barton, L.C. Hannah, Identification and molecular characterization of shrunken-2 cDNA clones of maize, Plant Cell, 2 (1990) 581–588.

[8] M.A. Ballicora, J.R. Dubay, C.H. Devillers, J. Preiss, Resurrecting the ancestral enzymatic role of a modulatory subunit, J Biol Chem, 280 (2005) 10189–10195.

[9] P.A. Nakata, T.W. Greene, J.M. Anderson, B.J. Smith-White, T.W. Okita, J. Preiss, Comparison of the primary sequences of two potato tuber ADP-glucose pyrophosphorylase subunits, Plant Mol Biology, 17 (1991) 1089–1093.

[10] T.W. Okita, P.A. Nakata, J.M. Anderson, J. Sowokinos, M. Morell, J. Preiss, The Subunit Structure of Potato Tuber ADPglucose Pyrophosphorylase, Plant physiology, 93 (1990) 785–790.

[11] M.A. Ballicora, M.J. Laughlin, Y. Fu, T.W. Okita, G.F. Barry, J. Preiss, Adenosine 5’-diphosphate-glucose pyrophosphorylase from potato tuber. Significance of the N terminus of the small subunit for catalytic properties and heat stability, Plant Physiol, 109 (1995) 245–251.

[12] P.R. Salamone, I.H. Kavakli, C.J. Slattery, T.W. Okita, Directed molecular evolution of ADP-glucose pyrophosphorylase, Proc Natl Acad Sci USA, 99 (2002) 1070–1075.

[13] A.A. Iglesias, G.F. Barry, C. Meyer, L. Bloksberg, P.A. Nakata, T. Greene, M.J. Laughlin, T.W. Okita, G.M. Kishore, J. Preiss, Expression of the potato tuber ADPglucose pyrophosphorylase in Escherichia coli, J Biol Chem, 268 (1993) 1081–1086.

[14] B. Cakir, A. Tuncel, A.R. Green, K. Koper, S.K. Hwang, T.W. Okita, C. Kang, Substrate binding properties of potato tuber ADP-glucose pyrophosphorylase as determined by isothermal titration calorimetry, FEBS Let, 589 (2015) 1444–1449.

[15] S.K. Hwang, Y. Nagai, D. Kim, T.W. Okita, Direct appraisal of the potato tuber ADP-glucose pyrophosphorylase large subunit in enzyme function by study of a novel mutant form, J Biol Chem, 283 (2008) 6640–6647.

[16] I.H. Kavakli, T.W. Greene, P.R. Salamone, S.B. Choi, T.W. Okita, Investigation of subunit function in ADP-glucose pyrophosphorylase, Biochem Biophys Res Commun, 281 (2001) 783–787.

[17] S.K. Hwang, S. Hamada, T.W. Okita, ATP binding site in the plant ADP-glucose pyrophosphorylase large subunit, FEBS Let, 580 (2006) 6741–6748.

[18] S.K. Hwang, S. Hamada, T.W. Okita, Catalytic implications of the higher plant ADP-glucose pyrophosphorylase large subunit, Phytochemistry, 68 (2007) 464–477.

[19] M.J. Giroux, J. Shaw, G. Barry, B.G. Cobb, T. Greene, T. Okita, L.C. Hannah, A single gene mutation that increases maize seed weight, Proc Natl Acad Sci USA, 93 (1996) 5824–5829.

[20] S.M. Lee, Y.H. Lee, H.U. Kim, S.C. Seo, S.J. Kwon, H.S. Cho, S.I. Kim, T. Okita, D. Kim, Characterization of the potato upreg1gene, encoding a mutated ADP-glucose pyrophosphorylase large subunit, in transformed rice, Plant Cell Tiss Org, 102 (2010) 171–179.

[21] Y. Obana, D. Omoto, C. Kato, K. Matsumoto, Y. Nagai, I.H. Kavakli, S. Hamada, G.E. Edwards, T.W. Okita, H. Matsui, H. Ito, Enhanced turnover of transitory starch by expression of up-regulated ADP-glucose pyrophosphorylases in Arabidopsis thaliana, Plant Sci, 170 (2006) 1–11.

[22] D.M. Stark, K.P. Timmerman, G.F. Barry, J. Preiss, G.M. Kishore, Regulation of the Amount of Starch in Plant-Tissues by Adp Glucose Pyrophosphorylase, Science, 258 (1992) 287–292.

[23] T.W. Greene, L.C. Hannah, Enhanced stability of maize endosperm ADP-glucose pyrophosphorylase is gained through mutants that alter subunit interactions, Proc Natl Acad Sci USA, 95 (1998) 13342–13347.

[24] P.L. Keeling, R. Banisadr, L. Barone, B.P. Wasserman, G.W. Singletary, Effect of Temperature on Enzymes in the Pathway of Starch Biosynthesis in Developing Wheat and Maize Grain, AustJ Plant Physiol, 21 (1994) 807–827.

[25] C.R.L. Linebarger, S.K. Boehlein, A.K. Sewell, J. Shaw, L.C. Hannah, Heat stability of maize endosperm ADP-glucose pyrophosphorylase is enhanced by insertion of a cysteine in the N terminus of the small subunit, Plant Physiol, 139 (2005) 1625–1634.

[26] S.K. Boehlein, J.R. Shaw, J.D. Stewart, L.C. Hannah, Heat stability and allosteric properties of the maize endosperm ADP-glucose pyrophosphorylase are intimately intertwined, Plant Physiol, 146 (2008) 289–299.

[27] A.B. Seferoglu, K. Koper, F.B. Can, G. Cevahir, I.H. Kavakli, Enhanced heterotetrameric assembly of potato ADP-glucose pyrophosphorylase using reverse genetics, Plant Cell Physiol, 55 (2014) 1473–1483.

[28] X.S. Jin, M.A. Ballicora, J. Preiss, J.H. Geiger, Crystal structure of potato tuber ADP-glucose pyrophosphorylase, Embo J, 24 (2005) 694–704.

[29] J.R. Cupp-Vickery, R.Y. Igarashi, M. Perez, M. Poland, C.R. Meyer, Structural analysis of ADP-glucose pyrophosphorylase from the bacterium Agrobacterium tumefaciens, Biochemistry, 47 (2008) 4439–4451.

[30] I. Baris, A. Tuncel, N. Ozber, O. Keskin, I.H. Kavakli, Investigation of the interaction between the large and small subunits of potato ADP-glucose pyrophosphorylase, PLoS Comp Biol, 5 (2009) e1000546.

[31] M. Danishuddin, R. Chatrath, R. Singh, Insights of interaction between small and large subunits of ADP-glucose pyrophosphorylase from bread wheat (Triticum aestivum L.), Bioinformation, 6 (2011) 144–148.

[32] C. Dawar, S. Jain, S. Kumar, Insight into the 3D structure of ADP-glucose pyrophosphorylase from rice (Oryza sativa L.), J Mol Model, 19 (2013) 3351–3367.

[33] N. Georgelis, J.R. Shaw, L.C. Hannah, Phylogenetic analysis of ADP-glucose pyrophosphorylase subunits reveals a role of subunit interfaces in the allosteric properties of the enzyme, Plant Physiol, 151 (2009) 67–77.

[34] A. Tuncel, I.H. Kavakli, O. Keskin, Insights into subunit interactions in the heterotetrameric structure of potato ADP-glucose pyrophosphorylase, Biophys J, 95 (2008) 3628–3639.

[35] P. Crevillen, M.A. Ballicora, A. Merida, J. Preiss, J.M. Romero, The different large subunit isoforms of Arabidopsis thaliana ADP-glucose pyrophosphorylase confer distinct kinetic and regulatory properties to the heterotetrameric enzyme, The J Biol Chem, 278 (2003) 28508–28515.

[36] T.W. Greene, L.C. Hannah, Maize endosperm ADP-glucose pyrophosphorylase SHRUNKEN2 and BRITTLE2 subunit interactions, Plant Cell, 10 (1998) 1295–1306.

[37] I.H. Kavakli, C. Kato, S.B. Choi, K.H. Kim, P.R. Salamone, H. Ito, T.W. Okita, Generation, characterization, and heterologous expression of wild-type and up-regulated forms of Arabidopsis thaliana leaf ADP-glucose pyrophosphorylase, Planta, 215 (2002) 430–439.

[38] I.H. Kavakli, J.S. Park, C.J. Slattery, P.R. Salamone, J. Frohlick, T.W. Okita, Analysis of allosteric effector binding sites of potato ADP-glucose pyrophosphorylase through reverse genetics, J Biol Chem, 276 (2001) 40834–40840.

[39] S.K. Hwang, P.R. Salamone, H. Kavakli, C.J. Slattery, T.W. Okita, Rapid purification of the potato ADP-glucose pyrophosphorylase by polyhistidine-mediated chromatography, Protein Express Purif, 38 (2004) 99–107.

[40] A.B. Seferoglu, I. Baris, H. Morgil, I. Tulum, S. Ozdas, G. Cevahir, I.H. Kavakli, Transcriptional regulation of the ADP-glucose pyrophosphorylase isoforms in the leaf and the stem under long and short photoperiod in lentil, Plant Sci, 205-206 (2013) 29–37.

[41] S.K. Boehlein, A.K. Sewell, J. Cross, J.D. Stewart, L.C. Hannah, Purification and characterization of adenosine diphosphate glucose pyrophosphorylase from maize/potato mosaics, Plant Physiol, 138 (2005) 1552–1562.

[42] J.C. Phillips, R. Braun, W. Wang, J. Gumbart, E. Tajkhorshid, E. Villa, C. Chipot, R.D. Skeel, L. Kale, K. Schulten, Scalable molecular dynamics with NAMD, J Comput Chem, 26 (2005) 1781–1802.

[43] R. Guerois, J. Nielsen, L. Serrano, Predicting changes in the stability of proteins and protein complexes: a study of more than 1000 mutations, J Mol Biol, 320 (2002) 369–387.

[44] C. Kiel, S. Wohlgemuth, F. Rousseau, J. Schymkowitz, J. Ferkinghoff-Borg, F. Wittinghofer, L. Serrano, Recognizing and Defining True Ras Binding Domains II: In Silico Prediction Based on Homology Modelling and Energy Calculations, J Mol Biol, 348(2005)759–775.

[45] C. Kiel, M. Foglierini, N. Kuemmmerer, P. Beltrao, L. Serrano, A genomewide Ras-effector interaction network., J Mol Biol, 370 (2007) 1020–1032.

[46] V. Kolsch, T. Seher, G. Fernandez-Ballester, L. Serrano, M. Leptin, Control of Drosophila Gastrulation by Apical Localization of Adherens Junctions and RhoGEF2, Science, 315 (2007) 384–386.

[47] E. Capriotti, P. Fariselli, I. Rossi, R. Casadio, A three-state prediction of single point mutations on protein stability changes, BMC Bioinformatics, 9 (2008) (Suppl 2):s6

[48] F. Pucci, R. Bourgeas, M. Rooman, Predicting protein thermal stability changes upon point mutations using statistical potentials: Introducing HoTMuSiC, Sci Rep, 6 (2016).

[49] N. Pokala, T. Handel, Energy functions for protein design: adjustment with protein-protein complex affinities, models for the unfolded state, and negative design of solubility and specificity, J Mol Biol, 347 (2005) 203–227.

[50] E. Capriotti, P. Fariselli, R. Casadio, I-Mutant2.0: predicting stability changes upon mutation from the protein sequence or structure, Nucleic Acids Res, 33 (2005) W306–W310.

[51] A. Benedix, C. Becker, B. de Groot, A. Caflisch, R. Böckmann, Predicting free energy changes using structural ensembles, Nature Methods, 6 (2009) 3–4.

[52] J. Schymkowitz, J. Borg, F. Stricher, R. Nys, F. Rousseau, L. Serrano, The FoldX web server: an online force field, Nucleic Acids Res, 33 (2005) W382–W388.

[53] C. Rohl, C. Strauss, K. Misura, D. Baker, Protein Structure Prediction Using ROSETTA, Methods Enzymol, 383 (2004) 66–93.

[54] Y. Dehouck, J. Kwasigroch, D. Gilis, M. Rooman, PoPMuSiC 2.1: a web server for the estimation of protein stability changes upon mutation and sequence optimality, BMC Bioinforma, 12 (2011) 151.

[55] J.M. Cross, M. Clancy, J.R. Shaw, T.W. Greene, R.R. Schmidt, T.W. Okita, L.C. Hannah, Both subunits of ADP-glucose pyrophosphorylase are regulatory, Plant Physiol, 135 (2004) 137–144.

[56] D. Kim, S.K. Hwang, T.W. Okita, Subunit interactions specify the allosteric regulatory properties of the potato tuber ADP-glucose pyrophosphorylase, Biochem Biophys Res Commun, 362 (2007) 301–306.

[57] S.K. Boehlein, J.R. Shaw, N. Georgelis, L.C. Hannah, Enhanced heat stability and kinetic parameters of maize endosperm ADPglucose pyrophosphorylase by alteration of phylogenetically identified amino acids, Arch Biochem Biophys, 543 (2014) 1–9.

